# Proteomic and Biological Profiling of Extracellular Vesicles from Alzheimer’s Disease Human Brain Tissues

**DOI:** 10.1101/733477

**Authors:** Satoshi Muraoka, Annina M. DeLeo, Manveen K. Sethi, Kayo Yukawa-Takamatsu, Zijian Yang, Jina Ko, John D. Hogan, Zhi Ruan, Yang You, Yuzhi Wang, Maria Medalla, Seiko Ikezu, Weiming Xia, Santi Gorantla, Howard E. Gendelman, David Issadore, Joseph Zaia, Tsuneya Ikezu

**Author notes:** These authors contributed equally to this manuscript (Satoshi Muraoka, Annina M. DeLeo, Manveen K. Sethi). **Correspondence author:** Tsuneya Ikezu, MD, PhD, Boston University School of Medicine, 72 East Concord St, L-606B, Boston, MA 02118, USA.

## Abstract

**Introduction:** Extracellular vesicles (EVs) from human Alzheimer’s disease (AD) biospecimens contain amyloid-β peptide (Aβ) and tau. While AD EVs are known to affect brain disease pathobiology, their biochemical and molecular characterizations remain ill defined.

**Methods:** EVs were isolated from the cortical grey matter of 20 AD and 18 control brains. Tau and Aβ levels were measured by immunoassay. Differentially expressed EV proteins were assessed by quantitative proteomics and machine learning.

**Results:** Levels of pS396 tau and Aβ were significantly elevated in AD EVs. High levels of neuron- and glia- specific factors are detected in control and AD EVs, respectively. Machine learning identified ANXA5, VGF, GPM6A and ACTZ in AD EV compared to controls. They distinguished AD EVs from controls in the test sets with 88% accuracy.

**Discussion:** In addition to Aβ and tau, ANXA5, VGF, GPM6A and ACTZ are new signature proteins in AD EVs.

## 1. Background

Alzheimer’s disease (AD) is the most common forms of dementia affecting nearly 50 million people worldwide. Neuropathologically, disease is characterized by amyloid plaques formed by extracellular aggregation of amyloid beta peptide (Aβ) and intracellular accumulation of neurofibrillary tangles (NFTs). These are formed in brain tissue by the hyperphosphorylated and misfolded microtuble-associated protein tau [1,2]. As AD progresses, Aβ and tau aggregates spread throughout the brain in a spatiotemporal manner [3,4]. Aβ deposition is most prominent in the frontal and temporal lobes, hippocampus and limbic system. Tau pathology, as classified by Braak and Braak, occurs in six histopathological stages. These correspond to tauopathy stages of AD. In stages I and II, NFTs appear in the entorhinal cortex and hippocampus,while in stage III and IV, higher densities extend beyond the entorhinal cortex and hippocampus to the neocortex. In the final V-VI stages, pathological tau deposits are present in throughout the hippocampus [3].

Extracellular vesicles (EVs), including exosomes (50-150nm), ectosomes/microvesicles (150-1000nm), and apoptotic bodies (1000-5000nm) are released from neurons, glia, and various other neural cells into the extracellular space [5–7]. These vesicles contain multiple forms of nucleic acids (microRNA, ncRNA, mRNA, DNA among others), lipids and proteins that are transferred from cell to cell, and found in blood, urine and cerebrosprinal fluid (CSF) [8,9]. In the central nervous system (CNS), it has been reported that AD-associated pathogenic proteins in brain EVs including tau and Aβ oligomers play important roles in AD pathogenesis [9–11][12,13].

Moreover, it has been reported that inhibition of EV synthesis reduced amyloid plaque deposition in the mouse model of AD, and stimulation of EV secretion increased intracellular transfer of prion protein in AD mouse models [14,15]. EVs are involved in the extracellular enzymatic degradation of Aβ and promote both Aβ aggregation and clearance by microglia [16,17]. Moreover, models of neuron-to-neuron transfer of tau seeds by EVs were reported [12,18–20]. Our own prior work demonstrated that microglia spread tau by EV secretion and that reducing EV synthesis significantly reduces tau propagation [10]. One mechanism centers around Bridging integrator 1 (BIN1), which is associated with the progression of tau pathology and observed by its abilities to alter tau clearance and by promoting the release of tau-enriched microglial EVs [21].

While EVs recovered from human and mouse brain tissues were examined by morphology, proteomics and RNA analyses [22–25], no comprehensive and quantitative proteomics database have yet been acquired for AD human brain tissues. Herein, we provide the first proteomic profiling of EVs isolated from postmortem AD and cognitively impaired control brain tissues. The analyses were combined with machine learning and quantitation of Aβ and tau species by epitope-specific ELISA. Machine learning identified and distinguished protein signatures of AD brain-derived EVs from controls with high degrees of accuracy.

## 2. Methods

### 2.1. Brain sample acquisitions

Two cohorts of brains were used in this study. The first cohort was obtained from the University of Nebraska Medical School (11 AD and 9 control) and the Greater Los Angeles Veteran’s Affairs Hospital (9 AD and 9 control) as part of NIH NeuroBioBank, which were matched for age and sex. The second cohort was obtained from the NIH NeuroBioBank (22 AD and 18 control), All were matched for age and sex. The Institutional Review Board at the University of Nebraska Medical School, Greater Los Angeles Veteran’s Affairs Hospital and the NIH NeuroBioBank approved the brain acquisitions provided by informed consent.

### 2.2. Purification of EVs from human brain samples

0.5 g of largely grey matter tissue from the frontal cortex of deceased AD or control cases were processed for EV extraction based on reported method with some modifications [23]. Briefly, frozen brain tissue was chopped on ice using a razor blade (# 12-640 Fischer Scientific) to generate 2-3mm^3^ sections. The sections were transferred to 3mL of Hibernate E solution containing 20 units of Papain (# LK003178 Worthington-biochemical corporation) in Earle’s Balanced Salt Solution (EBSS) (# 14155063 Gibco) and then incubated in a water bath at 37°C for 15 minutes by stirring once every 5 minutes. After incubation, the samples were immediately place on ice, and 6mL of ice-cold Hibernate E solution (# A1247601 Gibco). The dissociated brain tissue samples were gently homogenized (20 strokes) with a glass-Teflon homogenizer (# 89026-384 VWR). The homogenized tissue samples were filtered with 40 μm mesh filter (# 22-363-547 Fisher scientific). After filtration, the tissue samples were centrifuged at 300 × *g* for 10 min at 4°C (# 5720R Eppendorf). The supernatant from 300 × *g* was transferred to new 15mL tube and then centrifuged at 2,000 × *g* for 10 min at 4°C (# 5720R Eppendorf). The supernatant from 2,000 × *g* was transferred to 30mL conical tube and then centrifuged at 10,000 × *g* for 10 min at 4°C (# Avanti J-E JA25-50 Beckman Coulter). The supernatant from this spin was filtered through a 0.22-μm mesh filter (# SLGP033RS EMD Millipore) into new a polyallomer ultracentrifuge tube with 13.2mL capacity (# 331372 Beckman Coulter) and then was centrifuged at 100,000 × *g* for 70 min at 4°C (# Optima-XE SW41 Beckman Coulter). After ultracentrifugation, the supernatant was completely discarded, and the pellet was resuspended in 2mL of 0.475M of sucrose solution (# S5-3 Fisher science). All sucrose solutions were diluted in double-filtered PBS (dfPBS) with 0.22-μm pore size (# SLGP033RS EMD Millipore). The sucrose step gradient was created with six 2-mL steps starting from 2.0 M to 1.5M, 1.0M, 0.825M, 0.65M, and 0.475M (containing the resuspended pellet) in a polyallomer ultracentrifuge tube. Each layer was colored with commercially available food coloring to facilitate capture of the EV-rich interphase present between certain steps. The gradient was centrifuged at 200,000 × *g* for 20 h at 4°C (35,000 rpm with # Optima-XE SW41 Beckman Coulter), after centrifugation, the gradient was collected in 2mL fractions, except for the first and last fractions, which were 1mL each. Fraction III corresponded to the 1mL removed from the tube, and particles in this area had a buoyant density of approximately 1.08 g/cm^3^. The area between the second (0.65M) and third (0.825M) steps was collected and corresponded to fraction “V”, with a buoyant density of 1.10 - 1.12 g/cm^3^, while the interphase between the third and fourth steps corresponded to fraction “VI”, with a buoyant density of 1.12 - 1.15 g/cm^3^. The V and VI fractions were then diluted separately to a total volume of 12mL with dfPBS and centrifuged at 100,000 × *g* for 70 min at 4°C (# Optima-XE SW41 Beckman Coulter). The final pellet from each fraction was resuspended in 30 μL of dfPBS. For all biochemical analyses, these fractions were combined such that an equal volume of each fraction (V and VI) was used. For proteomics, an equal amount of total protein from each fraction (V and VI) was used.

### 2.3. Protein concentrations

The bicinchoninic acid (BCA) assay was used to determine protein concentration for each sample. The Pierce BCA protein assay kit was used (# 23225 Pierce). Due to the limited amount of sample, EVs were diluted 1:10 before loading into the assay, and a 10:80 ratio of sample to reaction components was used. All assays were allowed to incubate at 60°C for 30 min before protein concentration was read in a Biotek SynergyMix at 562nm. For all protein assays, raw readings were adjusted by dilution of the sample for the assay, and then by the final dilution of the sample and starting weight of the material. The average concentration was then taken for fractions V and VI.

### 2.4. Measurement of total tau (t-tau), phosphorylated tau (pT181 tau, pS199 tau, pS396 tau), amyloid beta (Aβ1-40, Aβ1-42) and Annexin V (ANXA5)

EVs were diluted 1:500 in TET buffer (50 mM Tris HCl pH 7.5, 2mM EDTA, 1% Triton X-100) supplemented with Pierce HALT inhibitor (# 78425 ThermoFisher) for t-tau and ANXA5 or TENT buffer (50mM Tris HCl pH 7.5, 2mM EDTA, 150mM NaCl, 1% Triton X-100) supplemented with phosphatase inhibitors (either addition of 0.5mM PMSF, 10mM NaF (# S7920 SIGMA), 200mM glycerol-2-phosphate (# 50020 SIGMA), and 1mM Na_3_VO_4_ (# S6508 SIGMA) or Pierce HALT inhibitor for pT181 tau, pS196 tau, and pS396 tau. Enzyme-linked immunosorbent assays (ELISAs) were performed to assess levels of t-tau, p-tau, and ANXA5. Commercially available kits from ThermoFisher (t-tau: # KHB0042, pT181: # KHO0631, pS199: # KHO0631, pS396: # KHO0631, ANXA5: # BMS252) were used according to manufacturer’s instructions. For Aβ1-40 and Aβ1-42, the levels were measured using the Single Molecule Counting (SMC®) Immunoassay Technology using commercially available kits from Millipore / Sigma (Aβ1-40: # 03-0145-00, Aβ1-42: # 03-0146-00).

### 2.5. Nanosight Tracking Analysis (NTA)

All samples were diluted in dfPBS at least 1:1000 or more to get particles within the target reading range for the Nanosight 300 machine (Malvern Panalytical Inc), which is 10-100 particles per frame. Using a syringe pump infusion system (Harvard Laboratories/Malvern), five 60-second videos were taken for each sample at 21°C. Analysis of particle counts was carried out in the Nanosight NTA 3.2 software (Malvern Panalytical Inc) with a detection threshold of 5. Particle counts were normalized for dilution on the machine, dilution of the final pellet, and starting material for EVs extraction. The average count was then taken for fractions V and VI.

### 2.6. Transmission electron microscopy (TEM)

The EV isolated from AD or control brain tissue were analyzed by TEM. 5µl of the EV sample was adsorbed for 1 min to a carbon-coated mesh grid (# CF400-CU EMS www.emsdiasum.com) that had been made hydrophilic by a 20-sec exposure to a glow discharge (25mA). Excess liquid was removed with a filter paper (# 1 Whatman), the grid was then floated briefly on a drop of water (to wash away phosphate or salt), blotted on a filer paper, and then stained with 0.75% uranyl formate (# 22451 EMS) for 30 sec. After removing the excess uranyl formate with a filter paper, the grids were examined and random fields were photographed using a JEOL 1200EX TEM with an AMT 2k CCD camera.

### 2.7. Mass spectrometry

#### 2.7.1. Sample preparation

The EV samples were mixed with 2,2,2-Trifluoroethanol (TFE) (# 75-89-8 Millipore) to a final concentration of 50%. The samples were sonicated for 2 min on an ice-water bath (VWR Scientific) and then incubated at 60°C for 2 h. After cool down, the 5mM dithiothreitol (# 3483-12-3 Sigma Aldrich) was added to the samples, and then reduced for 30 min at 60°C. Further, the samples were alkylated with 10mM iodoacetamide (# 163-2109 BioRad) for 30 min at room temperature in the dark. The samples were digested with mass spectrometry grade trypsin (# V5280 Promega) in 50mM ammonium bicarbonate (pH 7.5) for protein digestion (1:30 w/w trypsin-to-protein) for 16h at 37°C. The digested peptides were dried by vacuum centrifugation (# SPD1010 Speedvac system, Thermo Savant). The dried samples were resuspended in 2% acetonitrile/water/0.1% trifluoroacetic acid (TFA) and desalted using C-18 spin columns (# 89870 ThermoScientific); the cleaned peptides were eluted using 60% acetonitrile (ACN)/water/0.1% TFA, dried by vacuum centrifugation (# SPD1010 Speedvac system, Thermo Savant) and further resuspended in 1% ACN/water/0.1% formic acid (FA) and analyzed by nano-liquid chromatography and tandem mass-spectrometry (Nano-LC-MS/MS).

#### 2.7.2. Liquid chromatography (LC)-electrospray ionization (ESI) tandem mass-spectroscopy (MS/MS) Analysis

Nano-LC-MS/MS analysis was conducted by a Q-ExactiveHF mass spectrometer (Thermo-Fisher Scientific) equipped with a nano ultra-performance liquid chromatography (UPLC) (Water Technology) Peptides were trapped on a trapping column (180 µm × 20 mm) and separated on a reversed phased C-18 analytical column (BEH C18, 150 µm × 100 mm) (Waters technology). We loaded 1μL onto the column and separation was achieved using a 75 min gradient of 2 to 98% ACN in 0.1% FA at a flow rate of ~500nL/min. Data-dependent acquisition tandem MS was acquired in the positive ionization mode for the top 20 most abundant precursor ions. The scan sequence began with an MS1 spectrum (Orbitrap; resolution 60,000; mass range 300-2000m/z; automatic gain control (AGC) target 1 × 10^6^; maximum injection time 100 ms). MS2 analysis consisted of higher energy collisional dissociation (Orbitrap; resolution 15,000; AGC 1 × 10^5^; normalized collision energy (NCE) 30; maximum injection time 100 ms, dynamic exclusion time of 8 s).

#### 2.7.3. Sequence database

The raw LC-MS/MS data were converted into mZML format using ProteoWizard msConvert [26]. The data were searched using PeaksDB and PeaksPTM using Peaks Studio version 8.0 (Bioinformatics Solutions, Inc., Waterloo, ON, Canada) against the Uniprot/Swissprot database for Homo sapiens with a 0.1% false discovery rate (FDR) and at least two unique peptides. A 10-ppm error tolerance for the precursor (MS1) and 0.02 Da mass error tolerance for fragment ions (MS2) were specified. A maximum of 3 missed cleavages per peptide was allowed for the database search, permitting non-tryptic cleavage at one end. Trypsin was specified as the enzyme and carbamidomethylation as a fixed modification. A PeaksPTM search was queued after the peaksDB search, using advanced settings of a larger set of variable modifications, including hydroxylation P, oxidation M, hydroxylation K, hydroxylation-Hex K, hydroxylation-Hex-Hex K, HexNAc ST, HexHexNAc ST, phosphorylation STY, ubiquitination K, deamidation N, methoxy K, and nitrotyrosine Y. The final protein list generated was a combination of peaskDB and peaksPTM searches. The label-free quantification was achieved using PEAKS Studio Quantification-label-free module with a setting of mass error tolerance of 10 ppm and a retention time shift tolerance of 2.0 min. For data filtering for label-free peptide quantification following parameters were used: significance 15, fold change 1, quality 0, average area 1E4, charges from 1-10. The protein quantification following settings was used: significance 0, fold change 1, at least 2 unique peptides, significance method PEAKS.

### 2.8. Statistical analyses

Statistical analysis was conducted using IBM SPSS software ver.25 and GraphPad Prism6. Between-group comparisons were analyzed by Student’s t-test, nonparametric Mann-Whitney U, or one-way ANOVA followed by Bonferroni correction for multiple comparisons. The Gene Ontology of identified proteins were elucidated by DAVID Bioinformatics Resources 6.8. (https://david.ncifcrf.gov). The Venn diagram was generated using Venny_2.1 (http://bioinfogp.cnb.csic.es/tools/venny/).

### 2.9. Machine learning

The protein biomarkers to distinguish patients with Alzheimer’s from controls were selected using Least Absolute Shrinkage and Selection Operator (LASSO) on the proteomics data from the training set (n = 21), where each patient’s true state is labeled. An ensemble machine learning classifier to evaluate the performance of the selected proteins was developed. The ensemble machine learning classifier consists of five individual machine learning algorithms to mitigate overfitting, including Linear Discriminant Analysis, Logisitic Regression, Naïve Bayes, Support Vector Machine, and K-Nearest-Neighbours [27,28]. The machine learning generated model’s performance was evaluated on a separate, user-blinded test set (n = 17).

## 3. Results

A total of 38 patient samples, composed of 20 AD (15M, 5F, Mean age: 75.0) and 18 control (14M, 4F, Mean age: 75.7) cases were used in study for EV biological and proteomics analyses. A total of 78 patient samples, composed of 42 AD (24M, 17F, Mean age: 81.0) and 36 control (22M, 15F, Mean age: 79.0) cases, were included for validation analysis **(Table 1)**. There were no statistical differences in the demographics between AD and controls with the exception of postmoterm intervals (PMI) of the validation set, which will be discussed in the validation study **(Figure 4E)**.

**Table 1.**
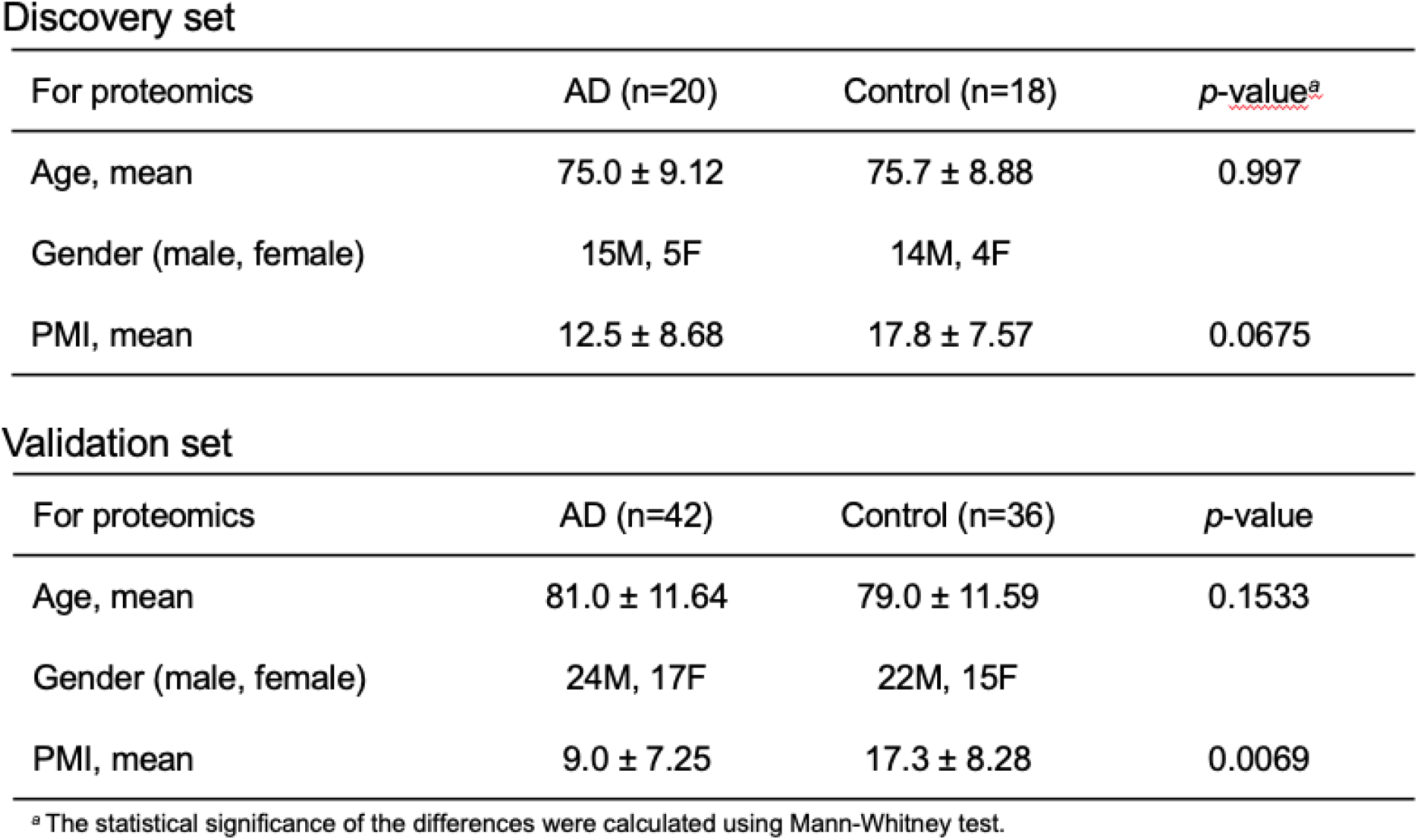
Patients information.

### 3.1. Experimental workflow

The experimental workflow is summarized in **Figure 1A**. The EVs were isolated from 20 AD and 18 sex- and age-matched cognitively unimpaired controls using the discontinuous sucrose gradient ultracentrifugation as previously described with modifications (see Materials and Methods) [23].

**Figure 1.**
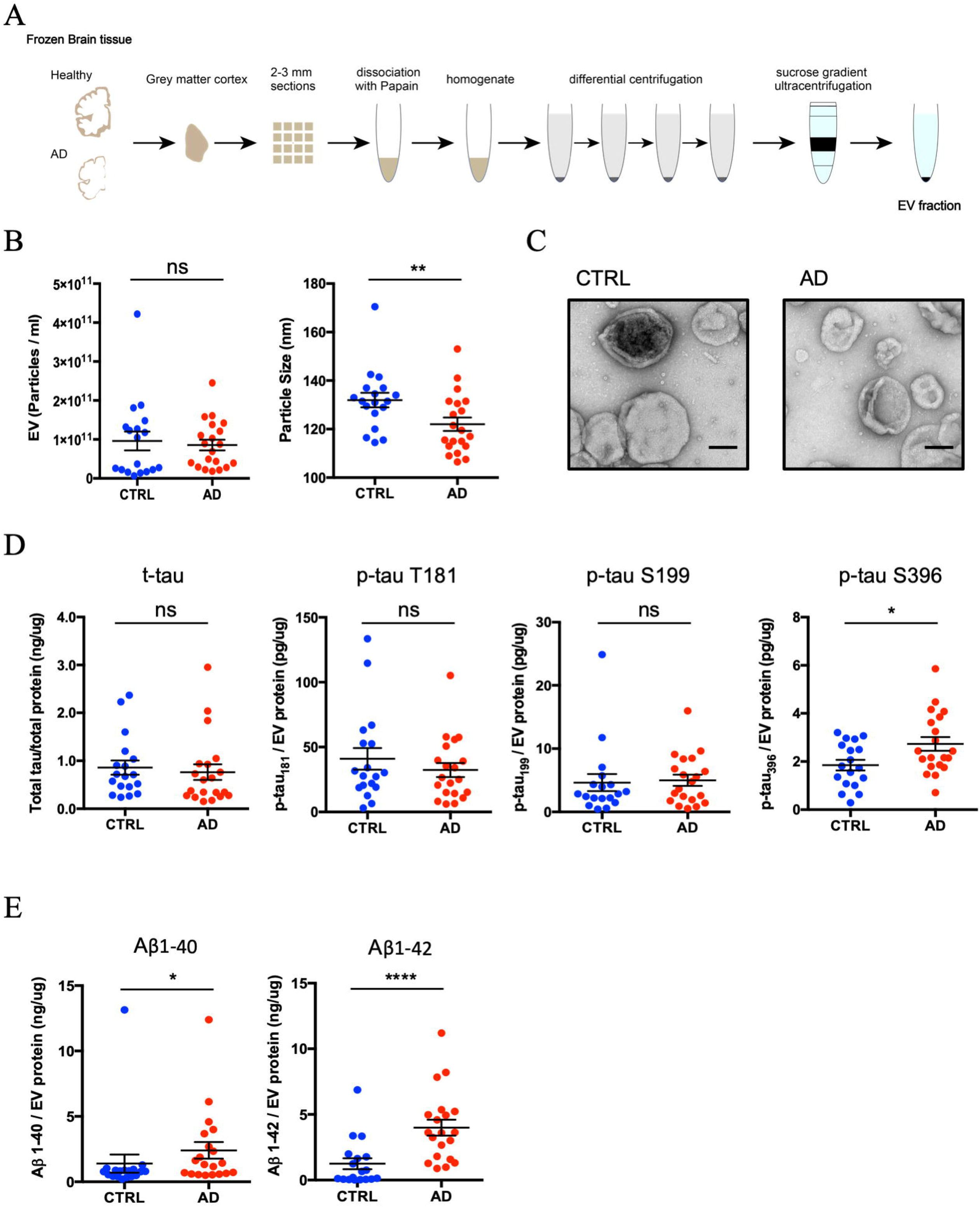
Biophysical and biochemical characterization of EVs isolated from AD and Control brain tissues: **A)** Schematic of extracellular vesicle isolation protocol from human frozen brain tissue. Frozen human gray matter cortex was chopped with a razor blade on ice to generate 2-3mm^3^ sections. The cut sections are dissociated in papain in Hibernate-E at 37°C for 15min. The tissue was homogenized with glass-Teflon homogenizer. After differential centrifugation, the EVs were purified by Sucrose step gradient Ultracentrifugation for 20 hours and then resuspended in double-filtered PBS. **B)** Left: Particle numbers of brain-derived EV fraction from control (CTRL) or AD by Nanoparticle tracking analysis. AD-brain derived EVs had no difference in number when adjusted for starting material than CTRL EVs (*p* = 0.6075 by Mann-Whitney test). Right: Particle size of brain-derived EV fraction. Average mode for particle size was smaller in AD brain compared with non-demented control brains (*p* = 0.0095 by Mann-Whitney test). **C)** TEM image of frozen human brain-derived EVs. EVs purified from AD, or non-demented CTRL brains were resuspended in double-filtered PBS and loaded onto formvar/carbon-coated copper mesh grids. The grids were then rinsed three times and negatively stained with 0.75% uranyl acetate. Scale bar is 100 nm. Left: CTRL, Right: AD. **D)** Comparison of total tau and tau phosphorylated at threonine 181, serine 199, and serine 396 in EVs. Tau is thought to be a major pathological factor in AD, especially phosphorylated, misfolded, and oligomerized tau. ELISA revealed that there is no difference in total tau to total protein (*p* = 0.398 by Mann-Whitney test). There was no difference in Tau phosphorylated at threonine181 to total tau (*p* = 0.7235) and at serine 199 to total tau (*p* = 0.4384). Tau phosphorylated at serine 396 to total tau was significantly different (*p* = 0.0375). **E)** Comparison of Amyloid beta 1-40 or 1-42 in EVs. ELISA revealed that there is a significant difference in amyloid beta 1-40 or 1-42 to total protein (*p* = 0.0259, *p* < 0.0001 by Mann-Whitney test).

### 3.2. Biochemical characterization of brain-derived EVs

To determine the purity of the EV preparations, NTA was deployed to determine the size of suspended particles based on their Brownian motion. Two cohorts of brain EVs (AD and control) and analyzed their size and number by NTA. The concentration of EVs derived from AD and control brains were not significantly different (*p* = 0.6075). The mode size distribution for EVs was significantly different and peaked at 122 nm for AD and 131 nm for controls (*p* = 0.0095) **(Figure 1B)**. The EVs isolated from frozen brain tissue showed cap-shaped morphology by transmission electron microscopy (TEM) **(Figure 1C)**. To characterize the AD-related proteins in brain-derived EVs, we measured the concentration of total tau (t-tau) and p-tau at threonine 181 (pT181 tau), serine 199 (pS199 tau) and serine 396 (pS396 tau) in lysed EVs by ELISA. The levels of t-tau, pT181 tau, and pS199 tau showed no significant differences between AD and controls (t-tau: *p* = 0.398, pT181 tau: *p* = 0.7235, and pS199 tau: *p* = 0.4384) **(Figure 1D)**. Conversely, pS396 tau was significantly increased in AD-brain derived EVs over controls (pS396 tau: *p* = 0.0375) **(Figure 1D)**. Moreover, we observed a significant increase in both Aβ1-40 and Aβ1-42 in AD-derived EVs over controls (Aβ1-40: *p* = 0.0021 and Aβ1-42: *p* < 0.0001) **(Figure 1E)**.

### 3.3. Proteomic profiling of brain-derived EVs

We performed a label-free Nano-LC-MS/MS analysis of 38 EV samples for proteomic profiling. Across both cohorts, a total of 1,088 proteins were identified with at least two unique peptides **(Figure 2A** and **Supplementary Table 1 and 2)**. There were 940 proteins identified in control EVs and 1,000 proteins identified in AD EVs. Among them, 852 proteins were detected in both groups, with 88 proteins unique to the controls and 148 proteins unique to the AD group **(Figure 2A)**. The common, AD-unique and control-unique proteins were tested for properties pertaining to the ‘cellular component’ and ‘pathway’ ontologies by Gene Ontology analysis in the Database for Annotation, Visualization and Integrated Discovery (DAVID). Among the 852 shared proteins, 60.9% were found to be included in the extracellular exosome ontology **(Figure 2B)**. The 148 proteins unique to the AD group were linked to mitochondria metabolism known to be dysfunctional in AD brain [29] **(Figure 2B)**. Interestingly, in pathway analysis by DAVID, neurodegenerative disorders, including AD, Parkinson’s and Huntington’s diseases were enriched in common and unique proteins **(Figure 2C)**. **Figure 2D** shows the peptides identified by Nano-LC-MS/MS, which covered 55.1% of tau (1-441), 9.1% of amyloid-beta precursor protein (APP 1-770), 74.3% of alpha-synuclein (SNCA 1-140) and 6.3% of apolipoprotein E (APOE 1-317). Post-translational modification (PTM) analysis detected six phosphorylation sites (pT181, pS198, pS199, pS202, pT231, and pS404) on tau (**Figure 2D**, red bold letters). Notably, Aβ sequence was identified in APP fragments. **Figure 2E** shows the AD pathway from KEGG pathway analysis based on 68 proteins identified in the AD group, which are designated with red stars. The list of AD pathway-related proteins is provided in **Supplementary Table 3**. Proteins known to play an important role in AD pathogenesis, such as APP, APOE, tau and NADH-ubiquinone oxidoreductase (Cx I-V), were all identified in the AD group, although they were not unique to this group.

**Figure 2.**
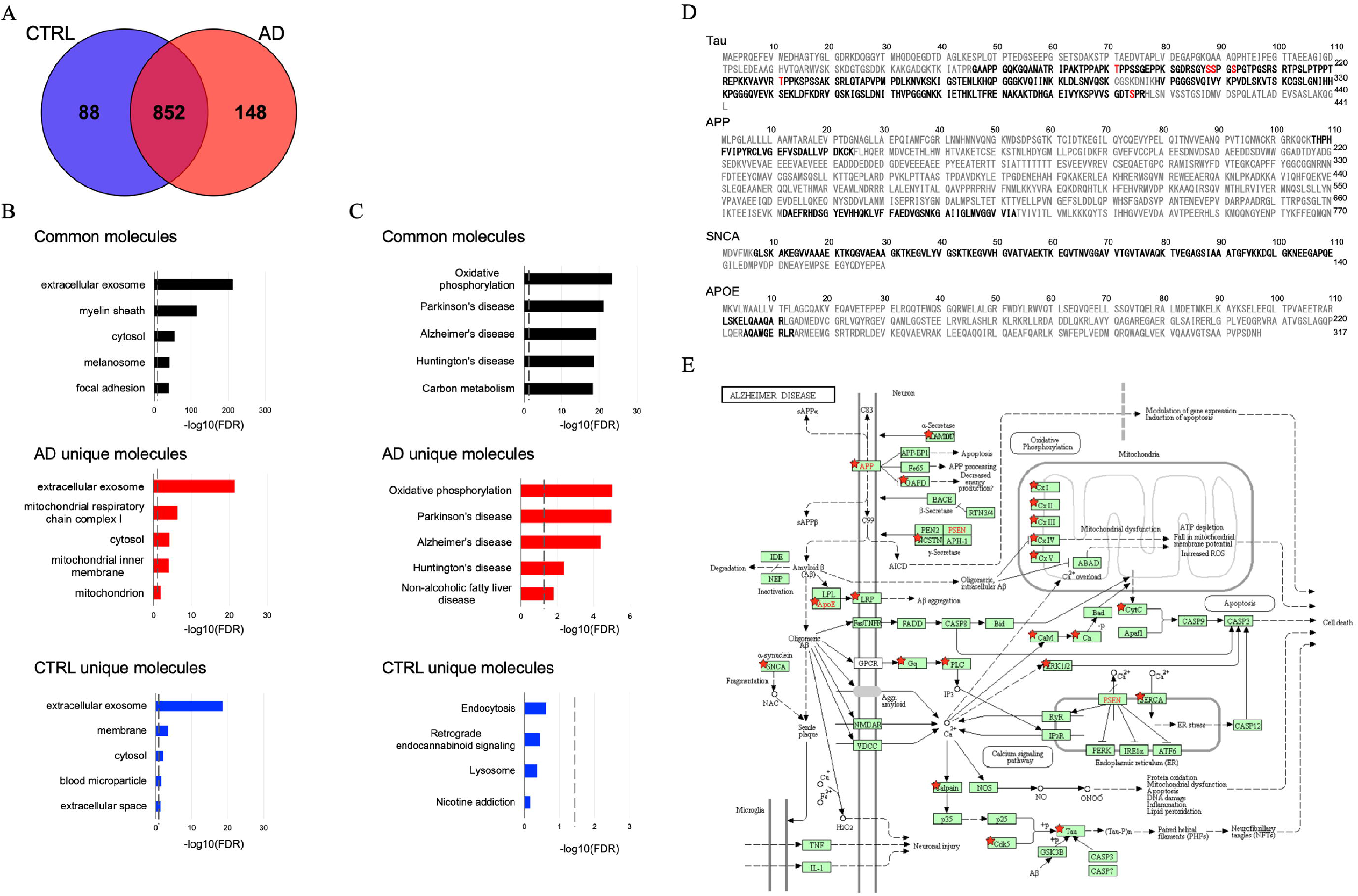
Proteomics profiling of EVs isolated from AD and CTRL brains: **A)** Venn diagram representing the number of EV proteins differentially identified in CTRL and AD. **B)** Gene Ontology (GO) analysis using DAVID Bioinformatics Resources 6.8. The GO term of Top5 Cellular Component with −log10 (FDR *p*-value). **(C)** The GO term of Top5 Pathway Ontology with −log10 (FDR *p*-value). **D)** Sequence coverage of identified tryptic fragment peptide from Alzheimer’s disease-related protein (Tau, APP, SNCA, APOE) by LC-MS/MS analysis. Identified peptides are shown in black bold. Identified phosphorylation sites are indicated in red bold. **E)** KEGG pathway of Alzheimer Disease. The 68 proteins identified in the AD group are highlighted by red stars.

### 3.4. Analysis of label-free quantitative proteomics comparison of AD and control brain-derived EVs

Label-free quantitative proteomics analysis was performed using PEAKS studio software. A total of 949 proteins were quantified (**Figure 3A**, **Supplementary Table 4**). The 934 quantified proteins were common between AD and control groups. Between these groups, 3 proteins were uniquely expressed in the controls, while 12 proteins were uniquely expressed in the AD group. The principal component analysis (PCA) showed a marginal separation of the two groups **(Figure 3B)**. **Figure 3C** shows the volcano plot of the common 289 proteins which were detected in more than 50% of the group (AD: n > 10 and controls: n > 9). Among these proteins, 15 proteins were significantly up-regulated and 3 proteins were significantly down-regulated in AD compared to the control group (as determined by p <0.05 and log2 fold change threshold of >1 or <−1) (**Figure 3C**, **and** **Table 2**). The expression levels of 18 molecules identified in the AD group relative to the control group are displayed in a heatmap **(Figure 3D)**. We next searched for brain cell-type specific molecules within the EV proteomics dataset using the mouse proteomics dataset as a reference [30]. The top 100 ranked cell type-specific molecules, which have at least 5-fold change in concentration in the cell type of interest over the other cell types, were tested with our EV proteomics dataset **(Figure 3E)**. The distribution of these markers indicate that in the human brain, neuron-derived EV represent 49% of EVs, while the other 50% of EVs are derived from glial cell types, including microglia, astrocytes and oligodendrocytes. Moreover, using label-free quantitative value, differences in the expression of cell type-specific marker molecules between AD and controls were seen **(Figure 3F)**. Interestingly, neuron-specific molecules were enriched in the control group (**Figure 3F**, **blue**), while glia-specific marker molecules were enriched in the AD group (**Figure 3F**, **red**). These results suggest that glia enhance their generation of EVs in AD brains, which may play a role in spreading AD pathologies.

**Figure 3.**
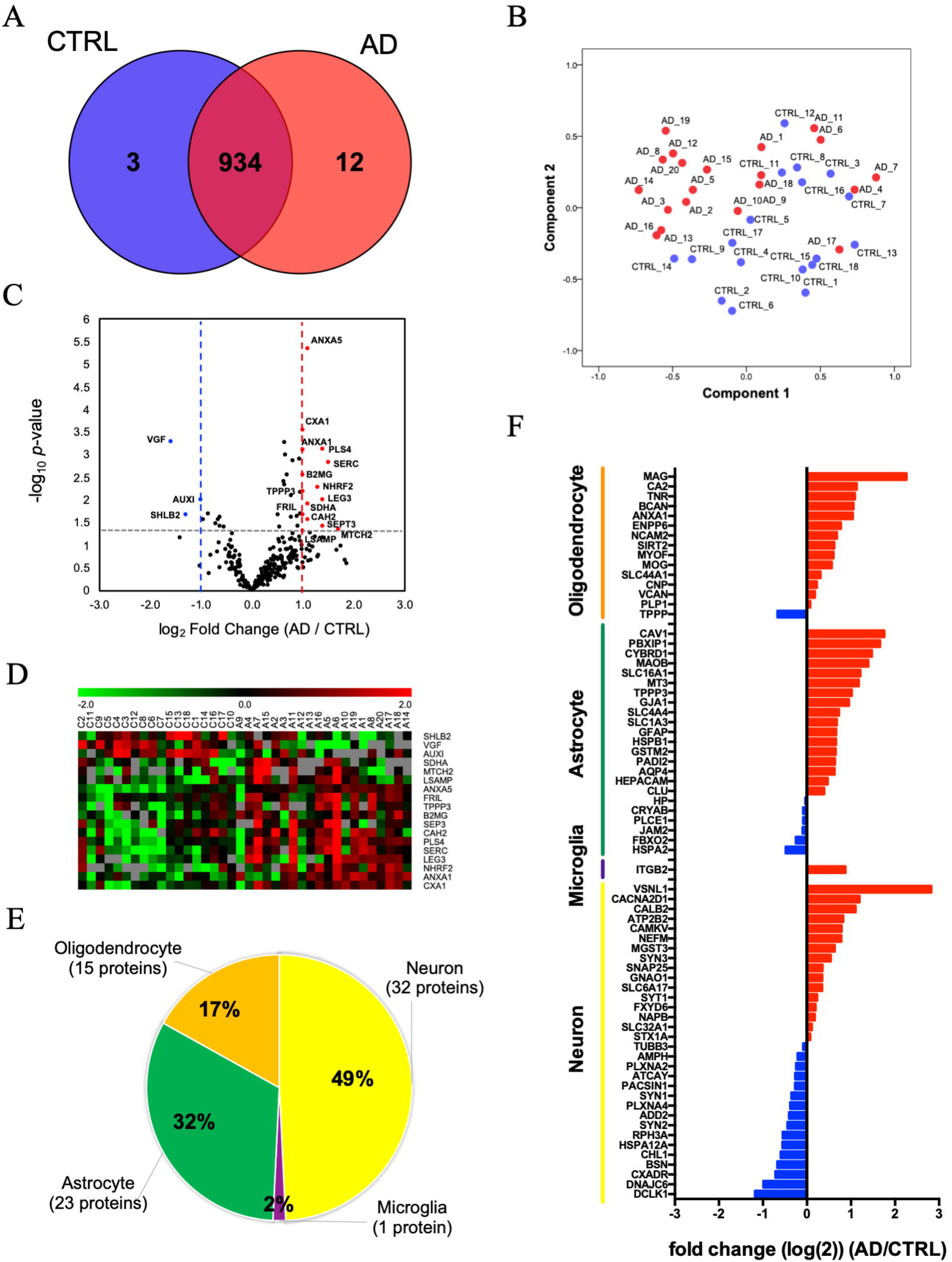
Label-free quantitative proteomics comparison of AD brain-derived EVs and CTRL brain-derived EVs: **A)** Venn diagram representing the number of EV proteins differentially expressed in CTRL and AD. **B)** The quantitative proteome profiles obtained by the label-free analysis were used to inform a principal component analysis (PCA). CTRL samples are defined by blue symbols, AD samples by red symbols. **C)** Volcano plot showing a degree of differential expression of EV proteins in AD compared with CTRL. The x-axis indicates log transformed fold change in expression. The y-axis shows log-transformed *p*-values. The grey dot line shows the 0.1 *p*-values and 1- or −1-fold change cutoff. **D)** Heat map representation of the up- and down-regulated proteins in AD. Red shows up-regulated proteins, and Green shows down-regulated proteins. **E)** Enrichment of brain cell type-specific markers in brain-derived EV proteins. Yellow: Neuron, Purple: Microglia, Green: Astrocytes, Orange: Oligodendrocytes. The parentheses show the number and percentage of identified cell type-specific protein. **F)** Comparison of the cell type-specific protein in AD brain-derived EV and CTRL EV. The red bar shows higher expression in AD. Blue bar indicates higher expression in CTRL.

**Table 2.**
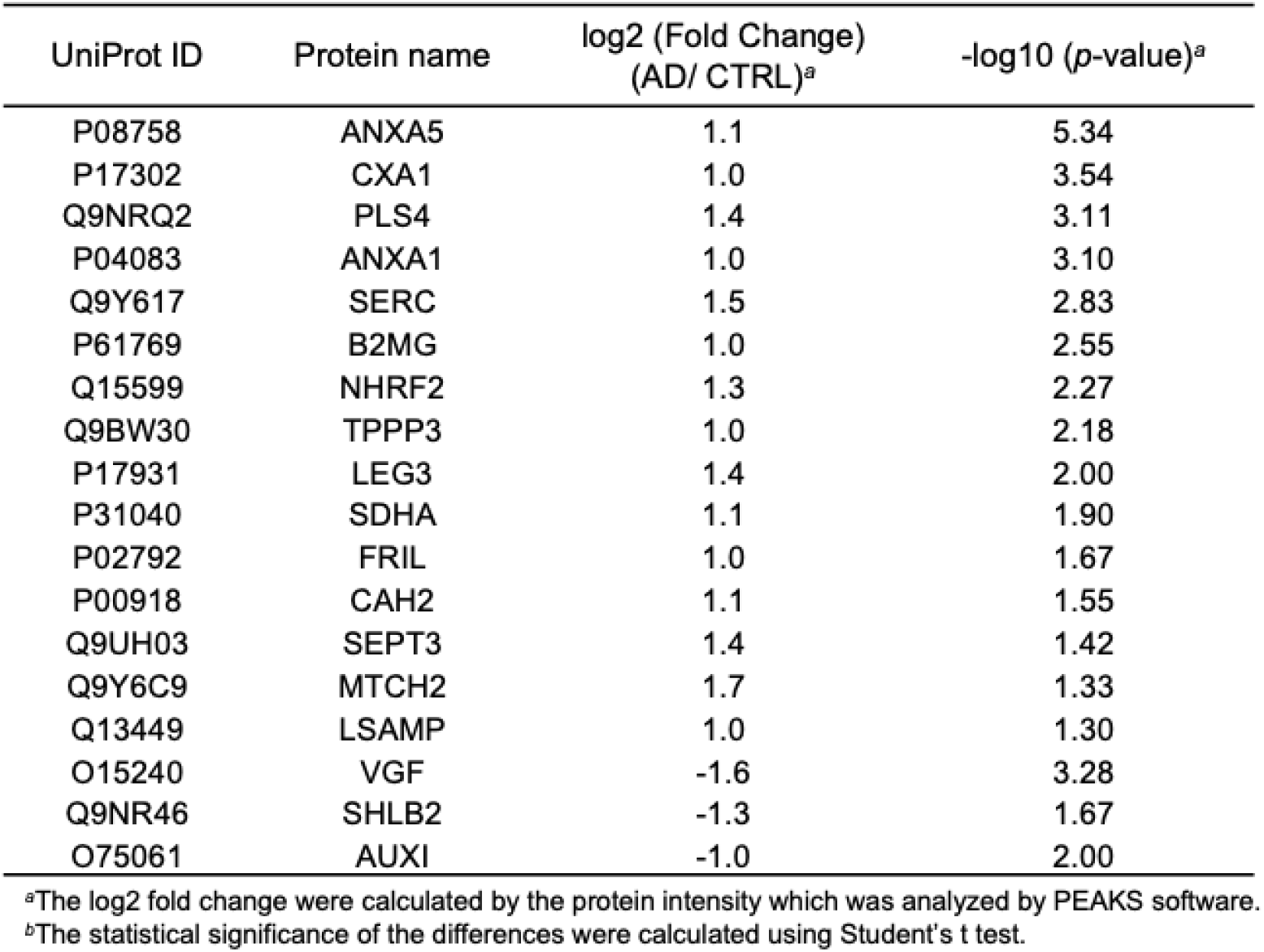
Up- and down- regulated protein in expression between AD and CTRL.

### 3.5. Machine learning to identify distinctive AD brain-derived EV proteins

To discover a combination of protein molecules that can accurately distinguish AD EVs from controls, the label-free quantitative proteomics dataset was analyzed using a machine learning method **(Figure 4A).** For this purpose, we split the proteomics dataset into an unblinded training subset (AD: n = 11; controls: n = 10) and a blinded testing subset (AD: n = 9; CTRL: n = 8). The ensemble machine learning model was built using only the data from the training subset, and then the accuracy of the diagnosis was determined using only the blinded testing set. We found that a panel that included annexin-A5 (ANXA5), NGF-induced growth factor (VGF), neuronal membrane glycoprotein M6-a (GPM6A), and alpha-centractin (ACTZ), selected by the LASSO algorithm, resulted in an area under the ROC curve (AUC) of 0.95 within the training set (**Figure 4B** and **Supplementary Table 5**). We then examined the accuracy in the independent blinded test set using the 4 proteins in the dotted green box **(Figure 4B)**. Using this model, we achieved an 88% accuracy (AUC = 0.97) in identifying AD patients using the panel that consisted of ANXA5, VGF, GPM6A, and ACTZ **(Figure 4C).** Further, we ran two control experiments that randomly selected 4 proteins from a total possible 949 proteins to form the diagnosis panel (repeated 20 times, AUC = 0.58, Accuracy = 55%) and shuffling the true labels of the subjects within the training set (AUC = 0.47, Accuracy = 48%). The control’s AUC was significantly worse than using the 4-protein panel’s AUC (*P*<0.001) **(Figure 4C)**. **Figure 4D** shows the scatter plot of selected 4 proteins from our panel individually, which were each significantly differentially expressed between AD and control groups **(Supplementary Table 6)**. Next, we assessed the correlation of expression of these candidate molecules to the levels of phosphorylated tau (p-tau), Aβ1–40, and Aβ1–42 in EVs by Pearson’s correlation analysis. There was a significantly positive correlation between GPM6A and pS396 tau, and between GPM6A and Aβ1-42 levels (pS396 tau: r = 0.380, *p* = 0.019; Aβ1-42: r = 0.387, *p* = 0.016), and a significantly negative correlation between VGF and Aβ1-42 levels (r = −0.538, *p* = 0.002) **(Figure 4E)**. We also examined the first and second cohorts of AD and control cases to validate ANXA5 expression level by ELISA. ANXA5 expression level was significantly increased in AD brain-derived EVs compared to the control group (*p* = 0.0042, 42 AD and 36 control cases, **Figure 4F**). Although there was statistical difference in PMI between AD and control cases in the validation cohort (**Table 1**), Pearson’s correlation analysis of PMI and ANXA5 levels showed no significant correlation (r = −0.165, *p* = 0.149). Thus, this correlation is not due to the difference in PMI between the two groups. Interestingly, ANXA5 expression level shows a tendency to increase along with Braak stages in an AD-dependent manner **(Figure 4F)**. Therefore, ANXA5 is a potential EV molecule for both distinguishing AD and control EVs and as a surrogate marker for Braak stage.

**Figure 4.**
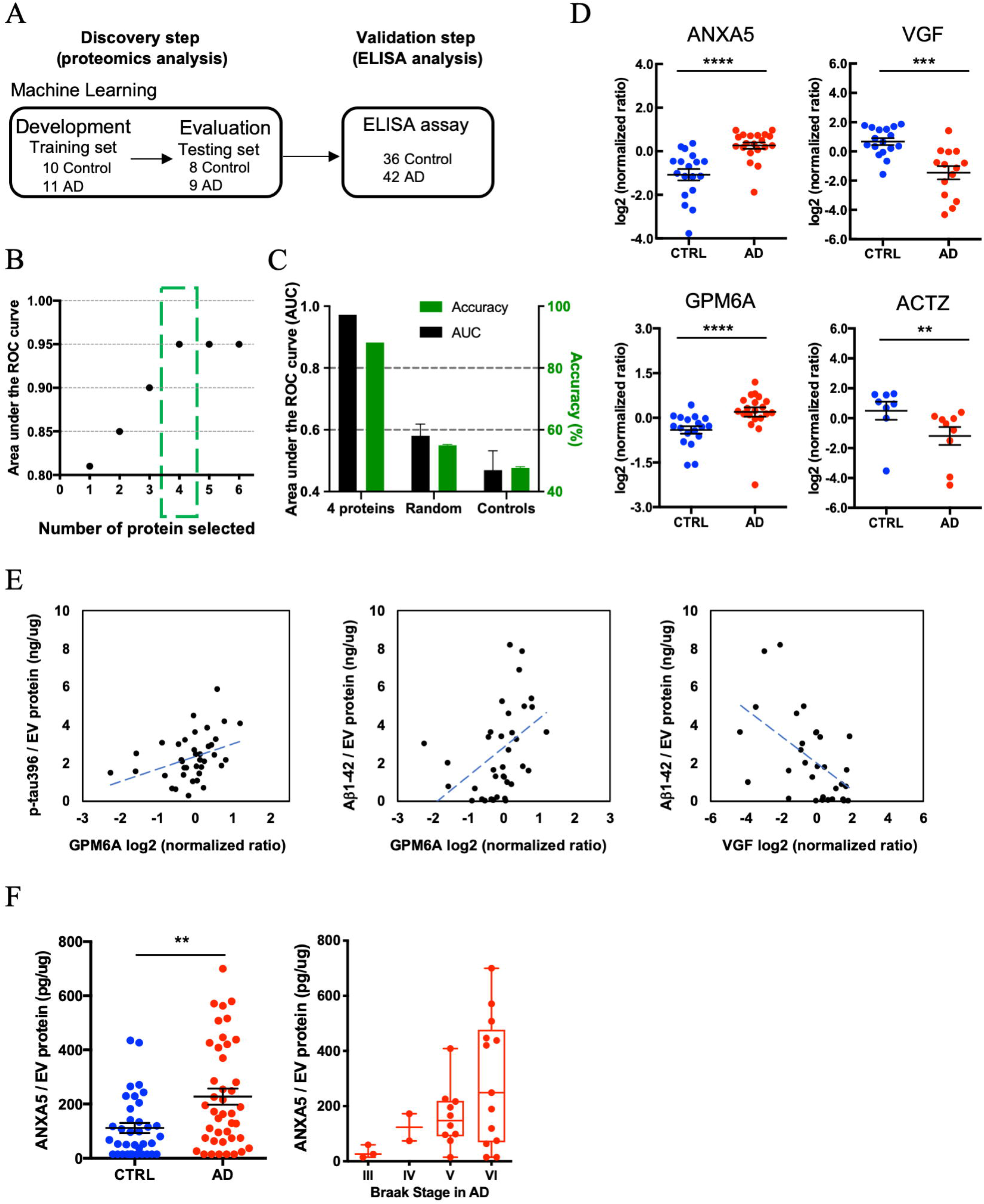
Machine learning model to identify AD brain-specific EV biomarker molecules: **A)** Workflow for Machine learning approach. The training set is fed into LDA to generate LDA vectors, which are applied to the blinded test set for classification. The predicted molecule are then validated from the validation cohort by a commercial ELISA. **B)** The best performing panel based on the area under the ROC curve using the Lasso algorithm was selected in the training set. The y-axis shows the area under the ROC curve. The x-axis indicates the number of proteins selected by the Lasso algorithm in the training set. The green box shows the selected protein with High AUC for the blinded test set. **C)** The Accuracy for 4 proteins with high AUC chose to generate the label of the test set, and note on their outcome to generate the prediction on the test set. Randomly selected control: Accuracy = 55%, AUC = 0.58., Shuffling control: Accuracy = 47.6%, AUC=0.47. **D)** A scatter plot of log2 (normalized ratio) as measured by proteomics per selected candidate protein in Machine Learning. (ANXA5: *p* = 4.58E-06, log2 fold change = 1.1, VGF: *p* = 5.30E-04, log2 fold change = −1.6, GPM6A: *p* = 5.37E-04, log2 fold change = 0.6, ACTZ: *p* = 4.44E-03, log2 fold change = −1.5). **E)** Scattered plot of candidate molecules and AD pathogenic molecule in brain-derived EVs. Left: GPM6A and pS396 tau (r = 0.380, *p* = 0.019). Center: GPM6A and Aβ1-42 (r = 0.387, *p* = 0.016). Right: VGF and Aβ1-42 (r = −0.538, *p* = 0.002). **F)** Left: Significant difference in total ANXA5 to total EV protein between AD and CTRL group by ELISA (*p* = 0.0042 by Mann-Whitney test) (AD = 42, CTRL = 36). Right: Braak stage-dependent increase in the ANXA5 expression level to total EV protein in AD dependent manner.

## 4. Discussion

The biophysical properties of EVs isolated from unfixed postmortem human brain tissues, quantitative analysis of tau and Aβ species, and label-free quantitative proteomic profiling by Nano-LC-MS/MS analyses were performed. pS396 tau, Aβ1-40, and Aβ1-42 levels were significantly increased in AD brain-derived EVs compared to controls. A total of 1,088 unique proteins from brain-derived EVs, were found to be enriched as extracellular exosomes molecules. We also quantified 949 proteins by label-free quantitative proteomic analysis, which were enriched in neuron-specific molecules in controls and glial cell type-specific molecules in the AD group. Differentially expressed EV proteins between AD and control brain samples included ANXA5, VGF, GPM6A and ACTZ as determined by machine learning approach. Using the validation cohort with the larger sample size, the increased protein level of ANXA5 in the AD group, as compared to controls was confirmed by ELISA.

When characterizing brain-derived EVs isolated from AD and control samples by tau-specific ELISA, the pS396 tau levels were found to be significantly increased in the AD group as compared to the control group. Previous reports indicate pT181 tau to be an early PTM associated with AD, pS199 tau modification is thought to promote tau accumulation, and pS396 tau modification is associated with tau seeding activity and aggregation [31,32]. The enrichment of pS396 in EVs derived from AD brain tissue may indicate their spread and aggregation of tau in the AD brain. We also observed tau fragments from the mid-region (156-406), which is inclusive of proline-rich domains (151-240), and microtubule binding repeat domains (244-369). Tau can be cleaved by various proteases including calpain-1 and 2 (at R230), thrombin, cathepsins, asparagine endopeptidase (at D255, N368), and caspase-2 (at D314) and −3 (at D421) [33–36]. However, many tau fragments identified in AD patients have not been well characterized, and the proteases responsible for their generation have not all been identified. In CSF, 25-35 kDa fragments of tau have been used as an early marker of AD; however, the protease responsible for this cleavage event is unknown [37,38]. A further detailed investigation is needed to determine if tau is truncated by proteases and modified by kinases, and are enriched in AD brain-derived EVs.

In the present study, we observed that the size of EVs derived from AD brain samples was smaller than from control EVs. We propose two possible factors for this finding. First, cholesterol level might be related to EV size; the CNS is known to contain 25% of the body’s cholesterol and accumulation of cholesterol is reported to be associated with AD [39]. High cholesterol levels increase Aβ in cellular and animal models of AD, and the chemical that inhibits cholesterol synthesis reduces Aβ in this model [39–43]. Moreover, it has been reported that small EVs contain significantly higher cholesterol than large EVs [40,41]. Therefore, AD brain cells that accumulate high cholesterol might secrete smaller EVs than controls. The second possibility is that the cell of origin determines EV size. Our study found that human brain-derived EVs were enriched in cell-type specific molecules corresponding to 50% neuronal and 50% glial cell origins. Moreover, the AD brain-derived EVs were enriched with more glial-specific molecules compared to the control EVs. Thus, there is the possibility that the difference in EV size between AD and control groups is due to an increase in smaller, glia-produced EVs among AD patients.

Herein, we report for the first time, the quantitative proteomic analysis of a large-scale brain-derived EV samples isolated from AD and control patients. A total of 59.5 % of the identified 1,088 proteins were part of the extracellular exosome ontology, and similar enrichment rate of EV-specific molecules was previously reported [22]. Interestingly, in AD EVs, glia-specific marker molecules were enriched, while neuron-specific molecules were enriched in controls. Thus, there is a possibility that glia-derived EVs act as a vehicle to transfer pathogenic proteins into neurons in AD brains, which are then spreading and propagating tau throughout the brain.

In summary, we have quantified tau and Aβ levels and investigated proteomic profiles in brain-derived EVs from AD, found enriched glia-specific molecules in AD brain-derived EVs, and identified ANXA5, VGF, GPM6A and ACTZ as potential candidate molecules for monitoring the progression of AD. Zhang *et al.* have recently reported that *ANXA5* is associated with familial late-onset AD by GWAS [44]. Further, Sohma *et al.* have reported a significantly higher plasma level of ANXA5 in AD patients than in control groups [45]. A number of previous studies in CSF and brain tissue have reported markedly lowered concentration of VGF in AD patient groups compared to controls [46–49]. GPM6A expression level has been shown to be downregulated in the hippocampus of transgenic mice modeling AD [50,51]. It has been reported that GPM6A with palmitoylation is involved in the clustering of lipid rafts, which are themselves enriched in sphingolipids and cholesterol [52]. Thus, GPM6A might be loaded into EVs with high cholesterol levels in AD. In our study, ANXA5 was detected in brain-derived EVs from one brain tissue cohort by commercial ELISA, but VGF, GPM6A, and ACTZ were undetectable. Thus, assays with higher sensitivity detection are needed for the quantification of these molecules. Finally, the combination of cell type-specific molecules from brain cells, including ANAX5, VGF, GPM6A or ACTZ may serve as potential biomarker candidate molecules in AD patient blood samples.

## Acknowledgments

The author thank Maria Ericsson (Electron Microscopy Facility, Harvard Medical School) for electron microscopic imaging services, participating laboratories of NIH NeuroBioBank for providing frozen human brain tissue specimens, and Li Wu (University of Nebraska Medical Center) for providing human brain tissue specimens.

## Author Contributions

S.M., A.M.D., and T.I. designed research; S.M., A.M.D., M.K.S., K.Y-T., S.A.D., Z.R., Y.Y., and Y.K.W. performed research; S.M., A.M.D., M.K.S., Z.Y., J.K., J.D.H., W.X., J.Z., and T.I. analyzed data; S.G., and H.E.G. provided brain samples; S.M., and T.I. wrote the paper; S.M., A.M.D., M.K.S., S.I., J.K., Z.Y., M.M., D.I., W.X., J.Z., and T.I. edited the paper.

## Supplementary Data

The Supplementary data related to this article can be found at:

